# Automated Discovery of Patterns in T-Cell Receptor Physicochemical Signatures

**DOI:** 10.1101/2025.07.07.663455

**Authors:** Zohreh Shams, Emma Bishop, Leo Mckee-Reid, Jessica Rumbelow

## Abstract

Accurately distinguishing antigen-reactive from non-reactive T-cell receptors (TCRs) is critical for advancing TCR-based immunotherapies and vaccines. Predicting antigen reactivity from physicochemical properties of the TCR sequence alone could enable rapid, low-cost identification of TCRs of interest, accelerating therapeutic discovery. In this paper, we use the *Discovery Engine*, a novel system for automated knowledge discovery from data, to classify published tumour antigen-reactive and non-reactive TCRs collected from cancer patients. Beyond classification, the *Discovery Engine* extracts interpretable combinatorial patterns (e.g., combinations of CDR3 length, net charge, and hydrophobicity) that predict whether a TCR is antigen-reactive. These patterns point to biologically meaningful features linked to tumour antigen recognition and could inform rational TCR design and prioritization. Notably, over half of the predictive patterns involve features from both the alpha and beta chains, highlighting the importance of considering both in assessing antigen specificity.

## 1 Introduction

T-cell receptors (TCRs) play a pivotal role in cancer immunotherapy by enabling T cells to specifically recognize and eliminate tumour cells through antigen recognition [1, 2]. Tumour-infiltrating lymphocytes (TILs), which are T cells found within the tumour microenvironment, have been harnessed for therapies with promising results [3, 4]. However, a significant challenge remains: not all TILs are tumour-reactive, as many may be bystanders or reactive to non-tumour antigens [5, 6]. Distinguishing tumour-reactive from non-reactive TILs is therefore critical for improving the efficacy of T cell-based cancer treatments.

Traditionally, identification of tumour-reactive TILs has relied on functional assays, surface marker profiling, or transcriptomic analyses, rather than on TCR sequence information alone [7, 8]. This is due in part to the complexity of TCR-antigen interactions and the fact that TCR sequence alone does not always predict antigen specificity [9]. As a result, most studies do not attempt to distinguish reactive from non-reactive T cells solely based on TCR sequence.

The TCR is a heterodimer composed of two polypeptide chains with multiple regions, including variable (V), joining (J), and complementarity-determining regions (CDRs), particularly the highly variable CDR3 loop, which is central to antigen recognition [2, 10]. Recent advances have made it possible to analyze the physicochemical properties of these individual regions and the TCR as a whole [11]. In this study, we leverage these properties to distinguish between tumour-reactive and non-reactive TILs using only TCR sequence information, providing a novel approach to T cell characterization in the context of cancer immunotherapy.

To classify antigen-reactive from non-reactive TCRs, we have turned to AI, which has shown great potential for scientific discovery (e.g., [12]). The successful examples however require (i) architecting a specific model for the task, which is time consuming, and (ii) potentially baking in some formalised domain expertise to help with the learning that can inevitably bias the model towards what is already known. More recently, Large Language Model (LLM) driven discovery pipelines [13, 14] have helped with the former problem, however, the latter is still a major issue in that it is impossible to judge what part of the discovery is in fact specifically related to a dataset in hand rather than the LLM’s knowledge based on its training data. This is on top of LLM shortcomings in mathematical tasks in general [15, 16].

A recent novel system called the Discovery Engine (Disco) [17] has addressed the limitations mentioned above by combining machine learning’s pattern recognition ability with state-of-the-art interpretability methods [18, 19] that aim at shedding light on decision making of machine learning models. This enables extracting human-understandable insights from complex data, called patterns, that would otherwise remain hidden. Disco is fully data-driven, fast, automated, and the patterns it extracts are reproducible.

We used Disco to classify antigen-reactive versus non-reactive TCRs, while also examining the features that differentiate them. We found that features related to the TCR alpha chain, such as CDR3 length and net charge, were informative for distinguishing between reactive and non-reactive TCRs. This is notable because the prevailing view has emphasized the greater variability and antigen-contact role of the beta chain, which is often considered more relevant for antigen specificity and binding [2, 10]. As a result, many studies that sequence only one TCR chain typically focus on the beta chain, leading to the alpha chain being comparatively less well-characterized in the literature [9, 8]. Our findings suggest that alpha chain features may play a more significant role in TCR reactivity than previously appreciated, highlighting the importance of comprehensive analysis of both chains.

In what follows we provide a brief overview of Disco, followed by the description of the dataset and the patterns extracted from it, along with their potential biological relevance.

## 2 The Discovery engine

Disco is composed of five main components:

- AutoPreprocess: This component automatically pre-processes the data. Operations done include imputation of missing values, duplication removal, elimination of correlated columns as well as handling of categorical and continuous variables.
- AutoML: The pre-processed data is modelled using an AutoML component, which includes several models in various categories: statistical (e.g., linear regression), tree-based (e.g., XGBoost [20]), kernel-based (e.g., SVM [21]) and deep learning models (e.g., autoencoders [22]).
- AutoInterp: A suite of state-of-the-art interpretability methods (e.g., [23, 24]) is applied to the trained models to elicit patterns in the data.
- AutoEval: This component distinguishes between patterns with strong empirical support in the data (validated patterns) that are statistically significant and those which demonstrate extrapolation from the data by the model (speculative patterns). It also applies LLMs to contextualize and assess the potential novelty of discovered patterns with respect to existing scientific literature.
- ReportGen: The final output of Disco is handled in this component, in the form of a PDF report outlining and evidencing the patterns found in the dataset, ranked by strength and novelty, with references to existing literature. It is also provided as a LaTeX document, well suited for sharing in academic contexts or for forming the basis of new publications.

## 3 The dataset

We compiled 2749 published paired-chain TCRs (2200 antigen-reactive, 549 non-reactive) collected from multiple cancer patients with a variety of malignancies. 2340 TCRs were sequenced from TILs in two patients [25], of which 2200 were non-reactive to tumour antigen and 140 were reactive. An additional 409 tumour-reactive TCRs were included from three curated datasets (Mc-PAS-TCR [26], IEDB [27], and NeoTCR [28]). The Peptides R package [29] was used in combination with a V gene-to-CDR lookup table from tcrdist3 [30] to infer physiochemical properties of the TCRs. This resulted in a dataset with 2749 rows and 18 columns/features, that are listed below.

Note that we concatenated CDR1 and CDR2 regions within the V gene and reported values derived from that as “cdrv”. The concatenated CDR1, CDR2, and CDR3 sequences make up “total_cdr” values. We also considered the V gene CDRs separately from CDR3 because the CDR3 region is so variable and might be important on its own.

- `a_total_cdr_len`: Total amino acid length of the alpha chain CDR1, CDR2, CDR2.5, and CDR3 regions
- `a_normalised_net_charge`: Net charge of the full alpha chain CDR1, CDR2, CDR2.5, and CDR3 amino acid sequence, normalised by its total length
- `a_hydrophobicity_gravy`: GRAVY (Grand Average of Hydropathy) score of the full alpha chain CDR1, CDR2, CDR2.5, and CDR3 sequence; a measure of average hydrophobicity
- `a_cdr3_total_cdr_len`: Total amino acid length of the alpha chain CDR3 region
- `a_cdr3_normalised_net_charge`: Net charge of the alpha chain CDR3 sequence, normalised by its length
- `a_cdr3_hydrophobicity_gravy`: GRAVY hydrophobicity index of the alpha chain CDR3 region
- `a_cdrv_total_cdr_len`: Total amino acid length of the alpha chain germline-encoded CDRs (CDR1, CDR2, and CDR2.5), excluding CDR3
- `a_cdrv_normalised_net_charge`: Net charge of the alpha chain CDR1, CDR2, and CDR2.5 sequence, normalised by its length
- `a_cdrv_hydrophobicity_gravy`: GRAVY hydrophobicity index of the alpha chain CDR1, CDR2, and CDR2.5 region
- `b_total_cdr_len`: Total amino acid length of the beta chain CDR1, CDR2, CDR2.5, and CDR3 regions
- `b_normalised_net_charge`: Net charge of the full beta chain CDR1, CDR2, CDR2.5, and CDR3 amino acid sequence, normalised by its total length
- `b_hydrophobicity_gravy`: GRAVY (Grand Average of Hydropathy) score of the full beta chain CDR1, CDR2, CDR2.5, and CDR3 sequence
- `b_cdr3_total_cdr_len`: Total amino acid length of the beta chain CDR3 region
- `b_cdr3_normalised_net_charge`: Net charge of the beta chain CDR3 sequence, normalised by its length
- `b_cdr3_hydrophobicity_gravy`: GRAVY hydrophobicity index of the beta chain CDR3 region
- `b_cdrv_total_cdr_len`: Total amino acid length of the beta chain germline-encoded CDRs (CDR1, CDR2, and CDR2.5), excluding CDR3
- `b_cdrv_normalised_net_charge`: Net charge of the beta chain CDR1, CDR2, and CDR2.5 sequence, normalised by its length
- `b_cdrv_hydrophobicity_gravy`: GRAVY hydrophobicity index of the beta chain CDR1, CDR2, and CDR2.5 region

## 4 The patterns discovered

In total Disco finds 29 patterns that are validated on the data. 7 focus on alpha chain properties only, 5 on beta chain properties only and more than half (17) on patterns combining features of alpha and beta chains. In what follows, we go through the discovered patterns, each of which is represented as a plot:

- The x-axis in each plot shows the pattern conditions, alongside the overall data. Conditions are listed above each plot and each indicates a certain range of values for a feature. The range of values is shown both in absolute terms and in terms of quantile in the data.
- The y-axis shows the proportion of the class the pattern is extracted for. In the context of the current dataset Reactive=1 refers to patterns related to antigen-specific TCRs, whereas, Reactive=0 refers to patterns related to non-specific TCRs.
- The notation above each bar in the plots shows the proportion of the class in y-axis (denoted as *µ*), the absolute number of data points satisfying the condition (denotes as *n*) and the significance of the condition (denotes as *p*, which stands for p-value).

### 4.1 Beta chain focused patterns

As discussed earlier, existing literature is mainly focused on the role of beta chain for antigen specificity and binding [2, 10]. Indeed Disco does find patterns that show this is the case in a proportion of the data. In the patterns in Figure 1 and 2, we see how beta chain CDR length, its hydrophobicity and its net charge play a role in increasing the reactivity, from the average rate of 19.9% in the overall data to 50% and 37.8%, respectively, under each pattern. The emphasis in both patterns is on low ranges of features appearing in the pattern. Interestingly, Disco finds a pattern, shown in Figure 3, that complements this finding very well. This pattern, unlike the previous two patterns, increases the proportion of non-reactive class from the average of 80.1% to 91.5%. It uses the same features as in Figure 2, however with emphasis on higher end of values for the features. Essentially contrasting patterns in Figure 2 and Figure 3 reveals that the combination of low values of beta chain CDR length and net charge increases the odds of reactivity, whereas, the combination of the higher values of these features increases the potential for non-reactivity.

**Figure 1:**
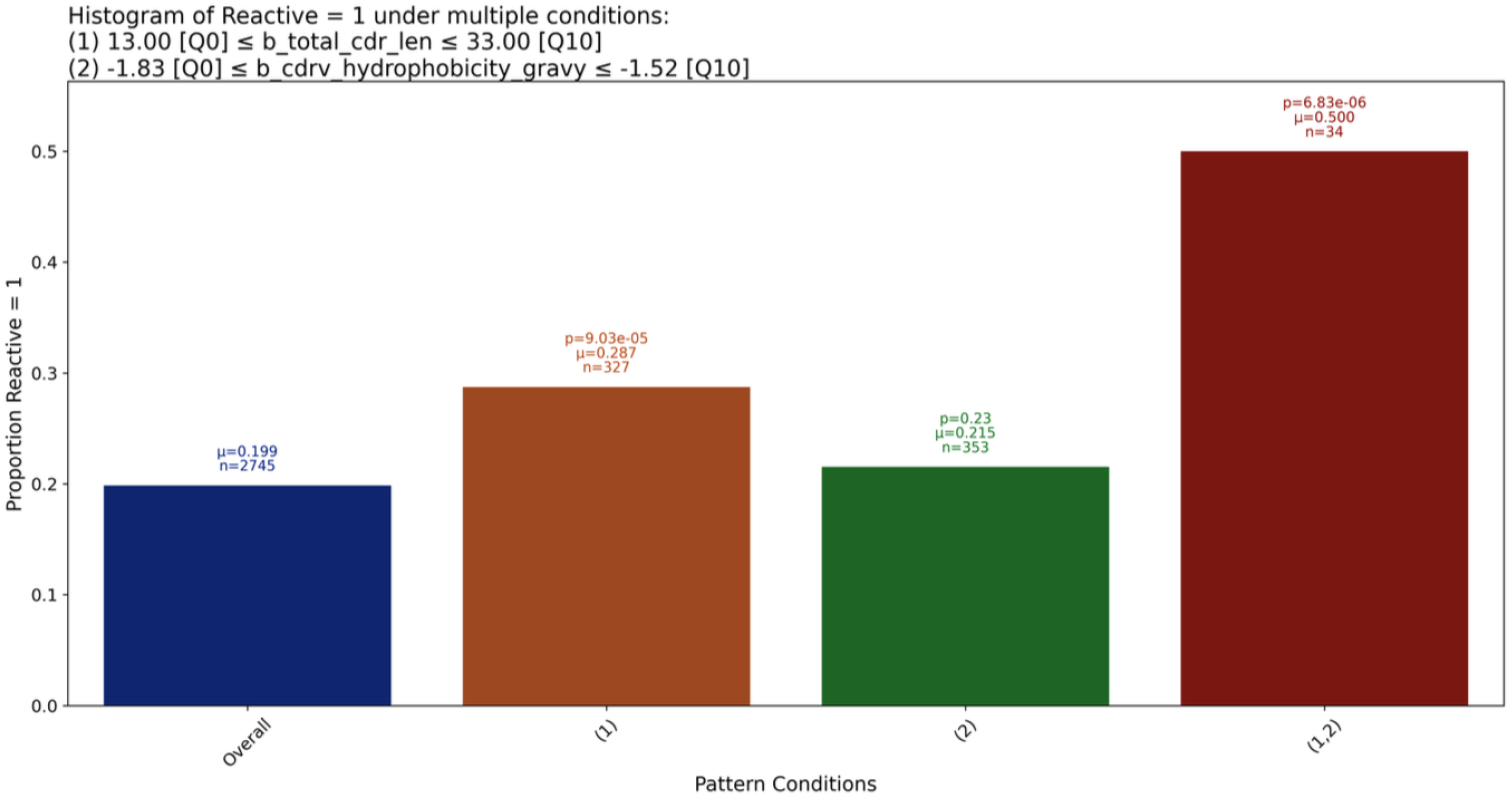
Low `b_cdrv_hydrophobicity_gravy` coupled with Short `b_total_cdr_len` increases the reactivity by *∼* 30%

**Figure 2:**
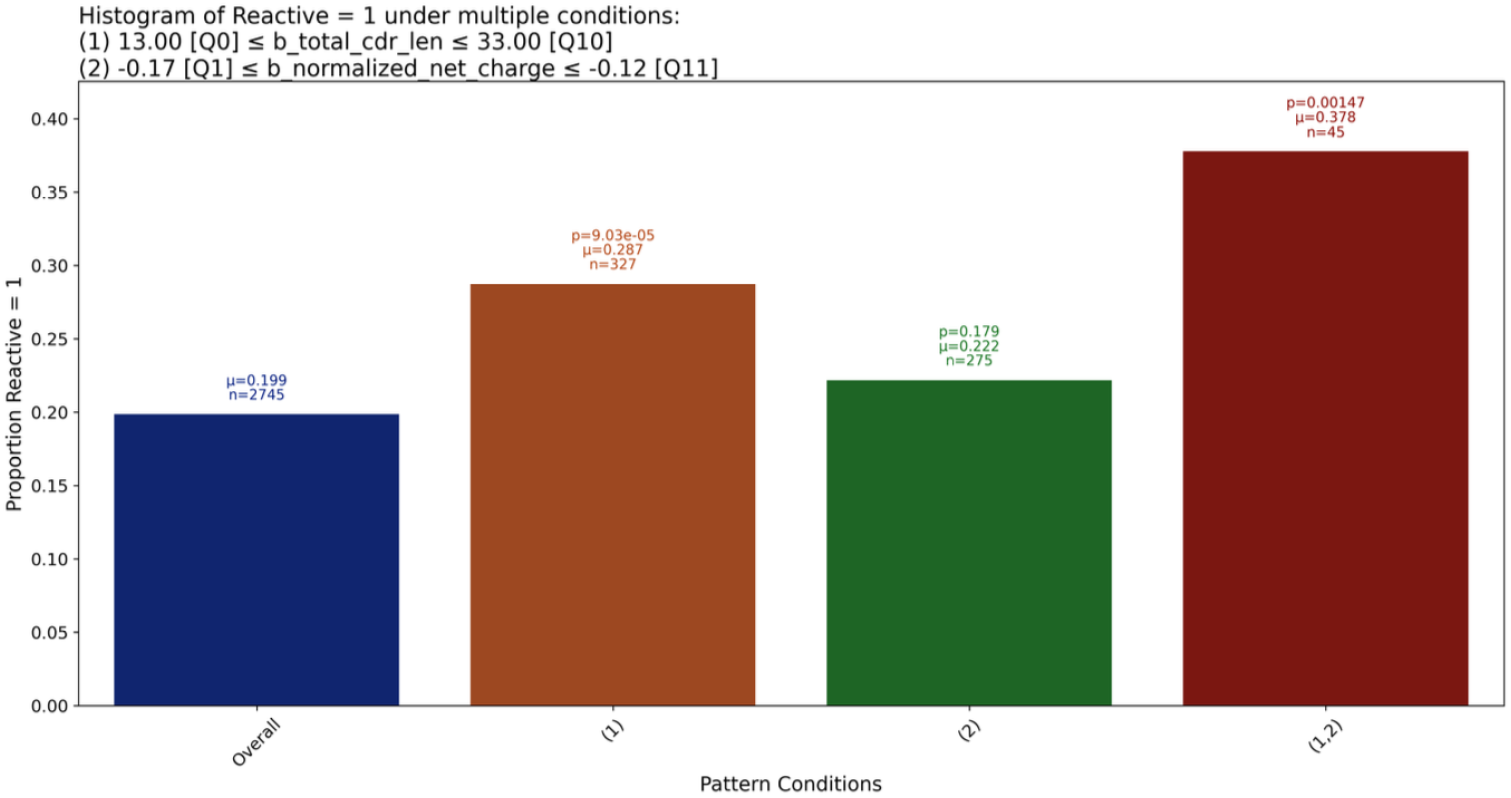
Low b_normalized_net_charge and short `b_total_cdr_len` increases the reactivity by *∼* 17%

**Figure 3:**
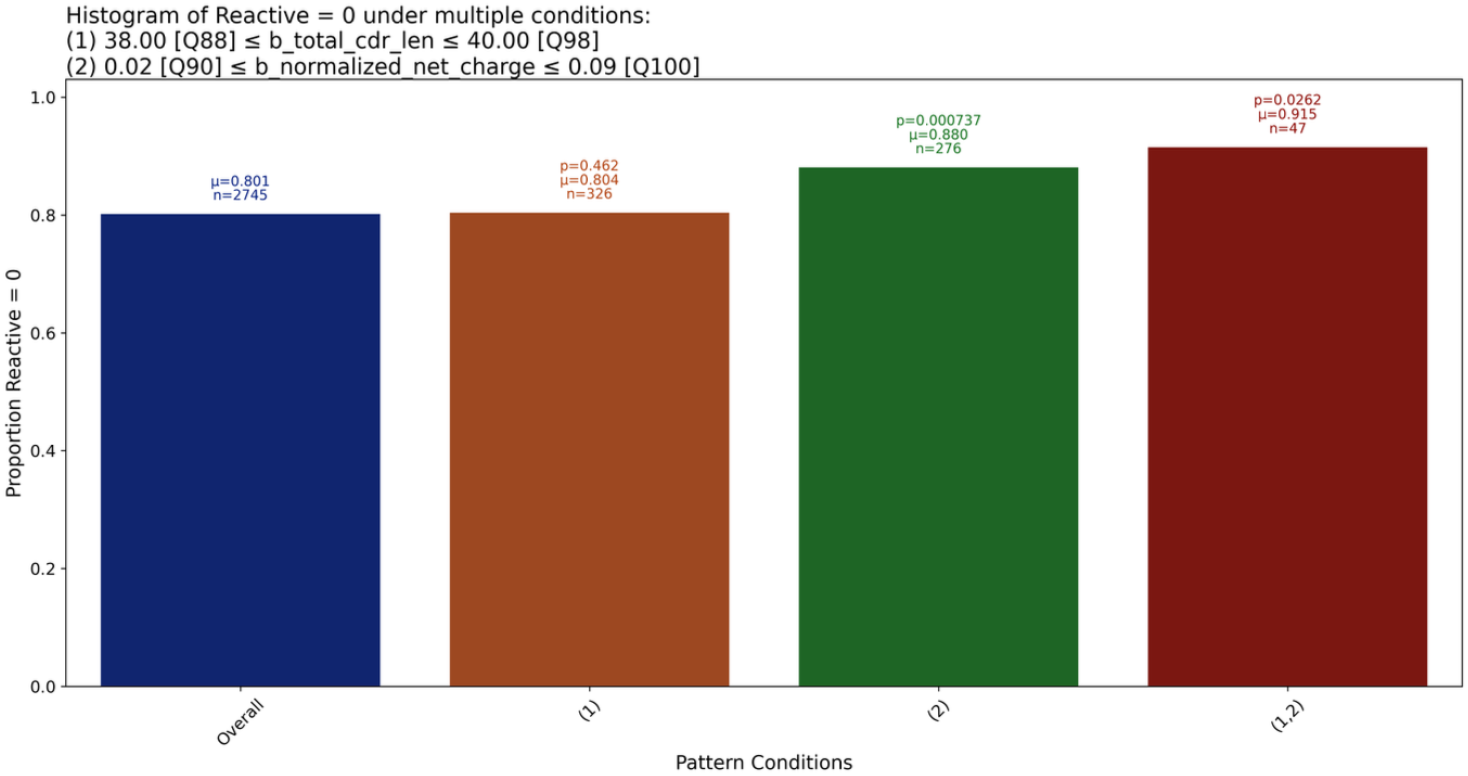
High b_normalized_net_charge and long `b_total_cdr_len` increases the non reactivity by *∼* 11%

### 4.2 Alpha chain focused patterns

Unlike the patterns in the previous section that are aligned with the literature, in that they are focused on the properties of beta chain, Disco finds significant patterns that are focused on alpha chain alone or a combination of alpha and beta chain properties. A notable feature in many of these patterns is `a_cdrv_total_cdr_len`, which is derived from the alpha chain V gene which encompasses the CDR1 and CDR2. All of these patterns are associated with reactive TCRs. In all of these cases, `a_cdrv_total_cdr_len` alone does not have much of an effect, but does when combined with other features (see patterns in Figures 4). Some researchers have shown a bias in the usage of alpha V gene TRAV12-2 in TCRs that respond to certain viral and cancer epitopes [31, 32]. In our data, only 44 out of 549 reactive TCRs use TRAV12-2, indicating that there could be a different V gene bias in our data or that the CDR1 and CDR2 lengths matter in their own right.

**Figure 4:**
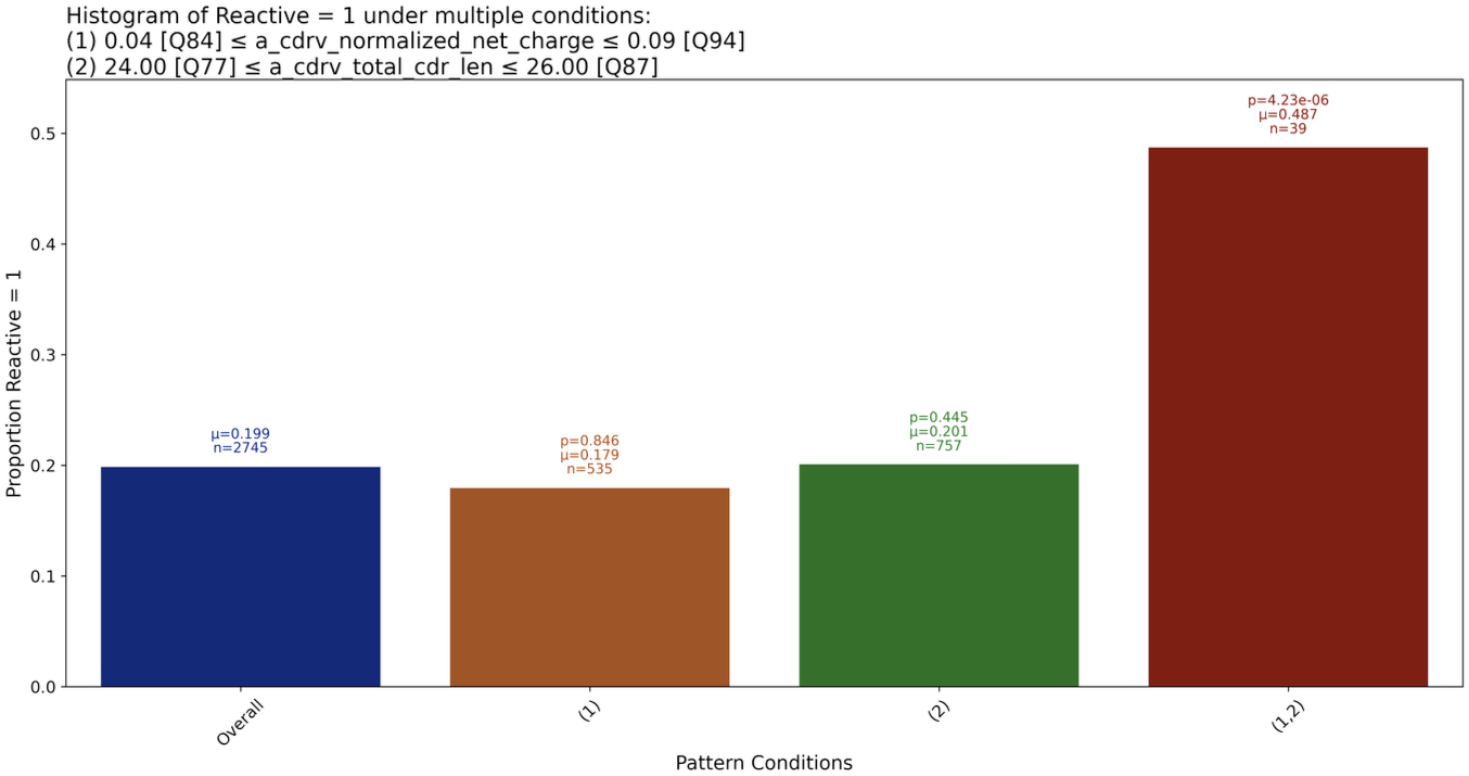
Alpha chain properties alone can increase reactivity by more than 25%

In addition to the above pattern, there is another interesting pattern (see Figure 5) that shows two features of the alpha chain which are not predictive alone but are together. These features are a_normalized_net_charge, derived from the entire alpha chain amino acid sequence, and `a_total_cdr_len`, derived from the alpha V gene and CDR3. This indicates that these features of the alpha chain could be more important for antigen recognition than previously thought but only in combination.

**Figure 5:**
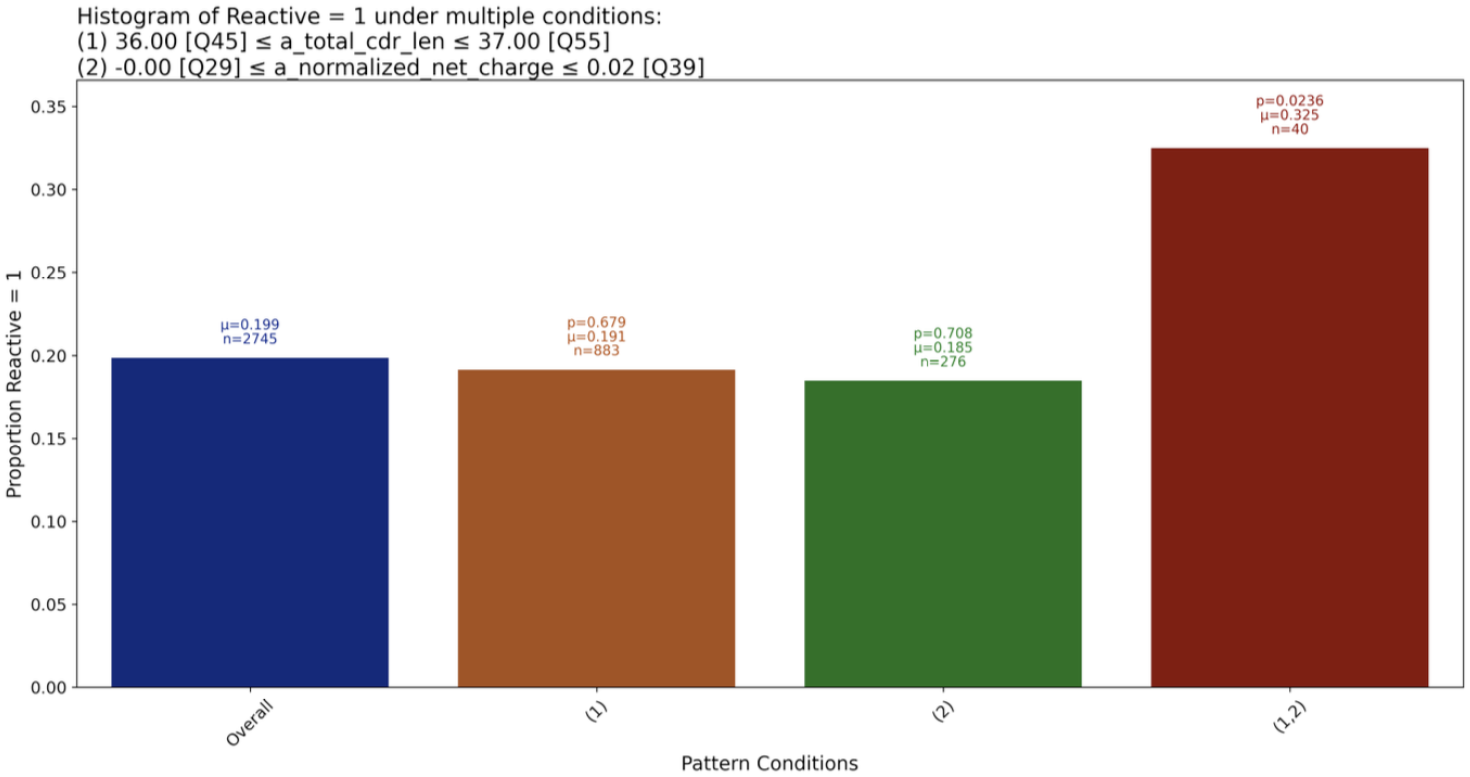
Combination of alpha chain properties can increase reactivity when individual components cannot

### 4.3 Patterns involving both chains

Whilst the focus of the previous two sections was on one chain only, the majority of the patterns (17 out of 29 validated ones) indeed include features of both the alpha and beta chains together to have a predictive effect. One of the most striking of these (pattern in Figure 6) is the combination of `b_cdr3_total_cdr_len`, the beta chain CDR3 length, and `a_cdrv_total_cdr_len`, derived from the alpha V gene. Alone, these features do not have a significant effect, but in combination they do.

**Figure 6:**
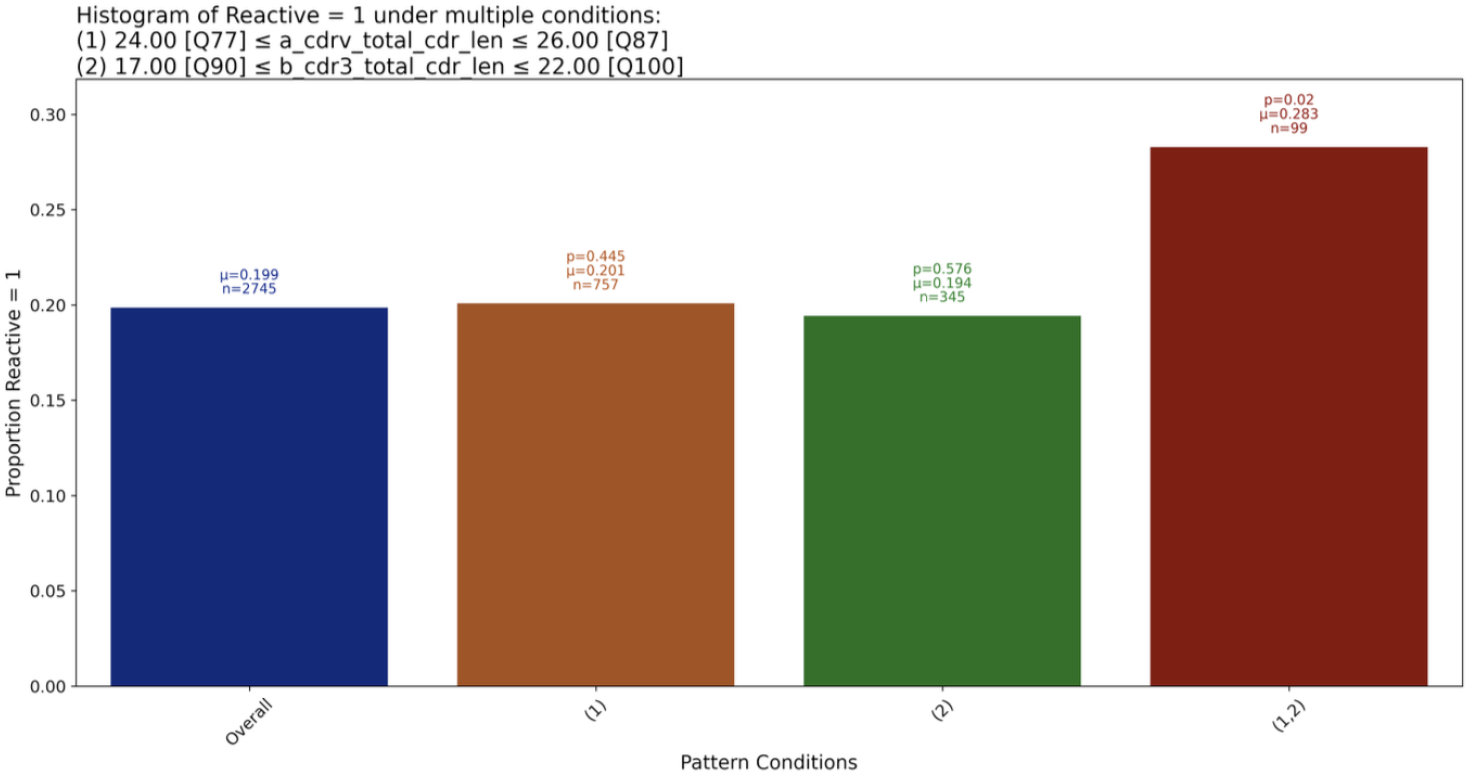
The impact of combination of alpha and beta chain properties

Similarly, Figure 7 shows a pattern involving both chains. The ranges for feature `b_cdr3_total_cdr_len` are identical to the previous patterns, however, the feature is paired up with high values of `a_hydrophobicity_gravy` and `b_total_cdr_len`. Whilst this pattern is supported by fewer data points, it increases reactivity more than the previous one by *∼*4%.

**Figure 7:**
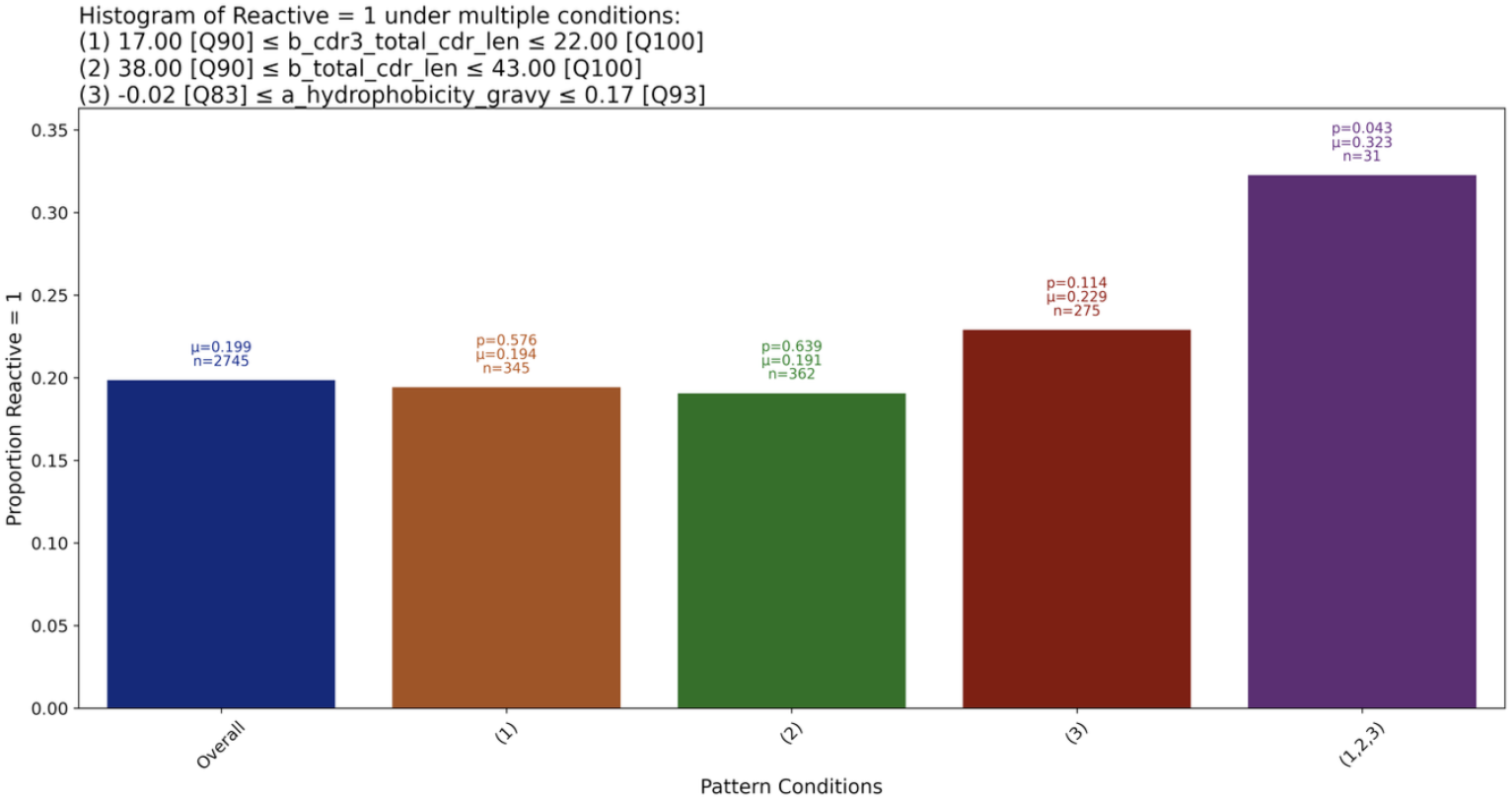
The impact of combination of alpha and beta chain properties

Finally, there is the pattern presented in Figure 8, which outperforms the previous two patterns in terms of the increase in reactivity. Surprisingly, in contrast to the pattern in Figure 7, this pattern is composed of very low values of b_total_cdr_le.

**Figure 8:**
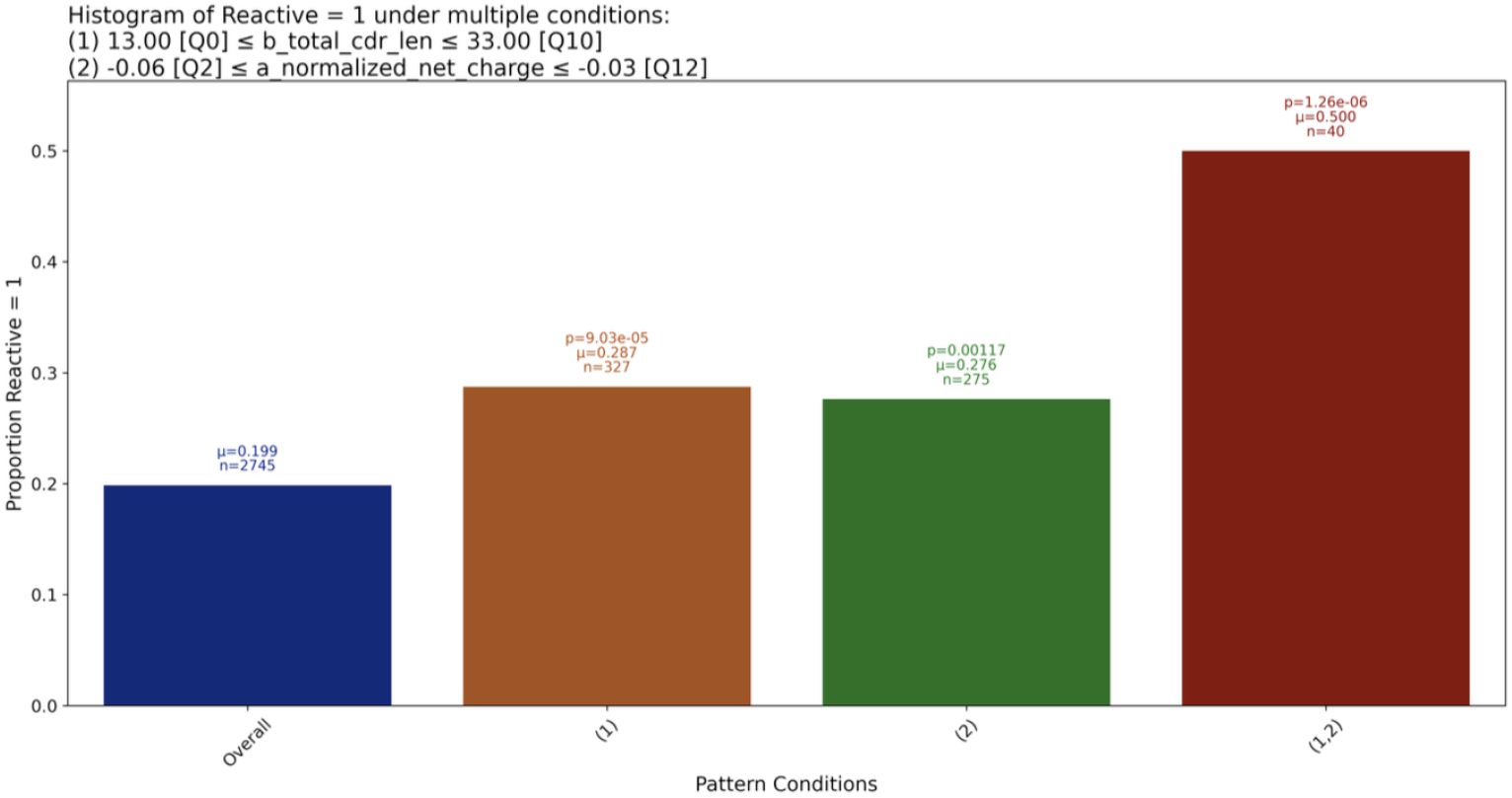
The impact of combination of alpha and beta chain properties

Indeed recent papers [33] indicate that TCR sequence alone is associated with T cell function, and they found this using paired-chain TCR data. This emphasizes the importance of studying the alpha and beta chains together. As it becomes cheaper and easier to generate paired-chain TCR data [34] we expect to be able to validate and reveal more patterns involving both chains.

## 5 Conclusion

In summary, our study demonstrates that features of the TCR alpha chain, such as CDR3 length and net charge, could be informative markers for distinguishing between tumour-reactive and non-reactive TCRs. These results challenge the prevailing focus on the beta chain and underscore the importance of analyzing both TCR chains when assessing antigen specificity and T cell reactivity. However, several limitations should be considered when interpreting our findings. Our non-reactive group included only two individuals, and the dataset was imbalanced, with a greater number of non-reactive TCRs compared to reactive ones. Additionally, the TCRs analyzed were sourced from a variety of contexts rather than a single tumour site, which may introduce heterogeneity and confounding factors. Despite these limitations, our findings provide new insights into the role of alpha chain features in TCR function and support the need for more comprehensive studies to further elucidate their contribution to tumour reactivity.

